# Mapping the space of protein binding sites with sequence-based protein language models

**DOI:** 10.1101/2024.07.24.604735

**Authors:** Tuğçe Oruç, Maria Kadukova, Thomas G. Davies, Marcel Verdonk, Carl Poelking

## Abstract

Binding sites are the key interfaces that determine a protein’s biological activity, and therefore common targets for therapeutic intervention. Techniques that help us detect, compare and contextualise binding sites are hence of immense interest to drug discovery. Here we present an approach that integrates protein language models with a 3D tesselation technique to derive rich and versatile representations of binding sites that combine functional, structural and evolutionary information with unprecedented detail. We demonstrate that the associated similarity metrics induce meaningful pocket clusterings by balancing local structure against global sequence effects. The resulting embeddings are shown to simplify a variety of downstream tasks: they help organise the “pocketome” in a way that efficiently contextualises new binding sites, construct performant druggability models, and define challenging train-test splits for believable benchmarking of pocket-centric machine-learning models.

## I. INTRODUCTION

Identifying protein-ligand binding sites and understanding their behaviour is crucial both for target discovery and structure-based molecular design [1–3]. Common objectives are to detect cryptic sub-pockets [4], identify allosteric sites or protein-protein interfaces [5], assess druggability [6], relate a site of interest to known pockets [7], transfer ligand information or hit matter across related sites [8], understand selectivity and pharmacophoric constraints [9], or elucidate the effect of mutations [10]. In practice, many of these analyses rely on techniques that efficiently search large databases such as the PDB for information on binding sites using appropriate filtering and ranking heuristics – i.e., *similarity metrics* [11, 12].

Many techniques that quantify pocket-pocket similarity have been proposed [13–23], with significant differences in scope, focus and speed. Typical is the use of both structural and physicochemical properties to describe residues, interactions and/or the pocket surface or shape. Some techniques are ligand-specific (i.e., they use a known ligand to bias the results [14–16]), others ligand-agnostic [17–22]. Faster approaches typically sacrifice sensitivity and accuracy for the ability to run queries interactively on the fly.

Despite this apparent abundance of approaches, we argue that more work on the topic is needed, and motivate this with the following three disparate observations:

- Transferable metrics that strike the “right” balance between structural, physicochemical, sequence and functional components remain a challenge.
- Protein-language models produce expressive vector embeddings that capture functional, evolutionary and structural relationships simultaneously and with low computational cost [24, 25].
- Pocket-centric machine-learning models (for example, for binding affinity or druggability prediction) are still often benchmarked with inappropriately debiased train-test splits that can produce unrealistic performance estimates [26].

The last point highlights that expressive similarity metrics that bridge multiple scales of information have applications beyond traditional target discovery and structure-based design. It is well established that debiased train-test splits are a prerequisite for believable benchmarks that enable fair comparisons while providing accurate estimates of a model’s prospective performance. Still, issues such as memorisation and poor transferability of deep-learning architectures remain persistent, as highlighted by the recent PoseBusters study [26]. And yet, despite this and other important efforts to establish best practices for validating deep-learning models, the proposed validation sets still often reduce to simple chronological splits that are prone to data leakage and use sequence identity cutoffs as a stand-in for more sophisticated 3D structural debiasing.

In this work we combine 3D structural processing with protein-language models to construct EPoCS – a transferable, robust, general-purpose similarity metric for protein binding sites. We show that EPoCS successfully captures information at multiple scales, from function, to sequence, to local structure, while enabling visual queries and on-the-fly context-matching across the pocketome [27]. By combining EPoCS with a hierarchical clustering approach, we debias train-test splits for improved benchmarking of machinelearning models with tunable sequences of progressively harder splits. We illustrate the approach for the prediction of protein druggability, highlighting how splits based on conventional homology cutoffs result in accidental data leakage, leading to less challenging validation sets than can be achieved with EPoCS.

The paper is organised as follows: Section II describes the formalism that integrates protein-language models with 3D tesselation techniques. Here we also study the relationship and correlation with existing similarity metrics. Section III focuses on the visualisation and key characteristics of EpoCS in organising the pocketome and modelling local and global pocket-pocket cross-similarity. Section IV describes our case study of applying EPoCS-derived train-test splits to druggability modelling.

## II. POCKET CROSS-SIMILARITY

This section describes how we combine protein-language models, specifically, ESM-2 [28], with a 3D tesselation approach to construct a multiscale metric for pocket cross-similarity. We refer to this metric as EPoCS (ESM-driven Pocket Cross-Similarity).

Protein Language Models (PLMs) have made dramatic progress over recent years, fuelled by the development of transformer-based architectures such as BERT [29]. They are typically trained using unsupervised or self-supervised protocols that task the model with reconstructing individual masked amino acids given an input sequence. PLMs have been applied successfully to a variety of problems, including the prediction of protein structures [28], mutation effects [30], antibody design [31] and binding-site detection [32]. The fact that PLMs are so versatile points to how information-rich their learnt embeddings are, and how their implicit biochemical, functional, evolutionary and chemogenomic information content is an emergent property of the pretext task that the models are subjected to during training.

Just how well these models integrate with structural applications is best demonstrated by ESMFold [28], a protein structure prediction model that substitutes AlphaFold2’s Evo-Former with an embedding layer derived from an ESM language model. However, even on its own, attention patterns of the ESM model have been shown to correlate well with realspace contacts of amino acids in the folded structure. This is an encouraging observation that justifies why ESM embeddings should also be a reasonable foundation for binding-site representations, where simultaneous descriptions of functional, structural and evolutionary factors should prove beneficial.

Figure 1 summarises the EPoCS approach that derives such representations from PLM embeddings: Given an input protein sequence, we use ESM-2 to generate residue embeddings for every position in the sequence. These embeddings are then mapped onto an experimental 3D crystallographic complex of the protein with a reference ligand. A radially truncated Voronoi tesselation around the ligand relative to the amino acids of the protein is used to identify the surface residues associated with the pocket volume. For non-liganded sites, a *pseudo*-ligand grown artificially into the pocket volume can serve as a substitute. Mean pooling of the embedding vectors over the surface residues then gives rise to the EPoCS embeddings vector. The Euclidean distance between embedding vectors is used to measure the distance between pairs of binding sites and thus serves as input for further processing, e.g., for hierarchical clustering and dimensionality reduction (see section III). Please refer to *Methods* for further details on the language embeddings, tesselation approach, clustering and 2D graph projection.

**Figure 1.**
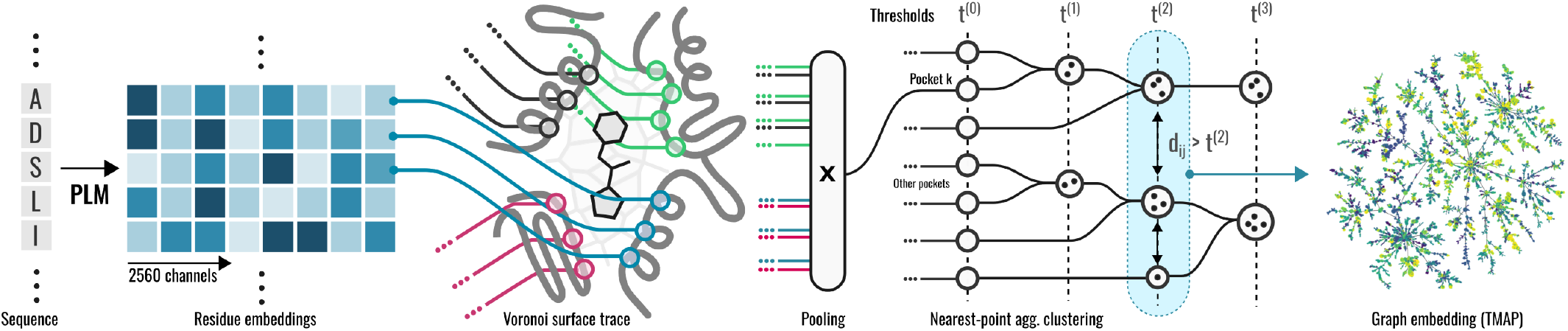
EPoCS approach for binding site representation and clustering. We use ESM-2 to generate embedding vectors for all residues of an input sequence. These embeddings are mapped onto the 3D structure of a protein-ligand complex. Radially truncated tesselation around a probe ligand identifies pocket interface residues of interest. Pooling of the embedding vectors over the interface results in the EPoCS embedding vector. The EPoCS metric is based on pairwise Euclidean distances between embedding vectors. Clusterings are obtained using an agglomerative strategy with strictly enforced minimum separations between clusters. We use tmap for visualisation of the metric space.

We argue that EPoCS is an intrinsically multiscale representation of a binding site due to its direct link to ESM embeddings via interface-based tesselation. To demonstrate this, we compare the metric space induced by EPoCS to two limiting cases: First, APoc (Alignment of Pockets) – a computationally demanding metric based on local residue features and local pocket alignments [17]; second, a simple sequence distance measure. We compile a random set of ∼3500 pockets from the PDB (here referred to as PDB-3.5k) and measure all pairwise distances between the pockets using these three different techniques. The resulting correlation plots are shown in Fig. 2a-c. Interestingly, EPoCS produces a good correlation with both APoc and sequence distance for pairs of structures that the two baseline metrics consider similar (see panels a and b) – whereas the correlation between the baselines themselves shows significantly more scatter (c). Furthermore, comparing the marginal distributions generated by the individual metrics, EPoCS produces a better contrast across its distance domain with a heavier tail in the distance distribution, as opposed to the sharply peaked distributions of the baselines. I.e., EPoCS captures both the local, regional and global similarity structure across the set, as will become even clearer when we visualise the metric space using projection techniques (section III).

**Figure 2.**
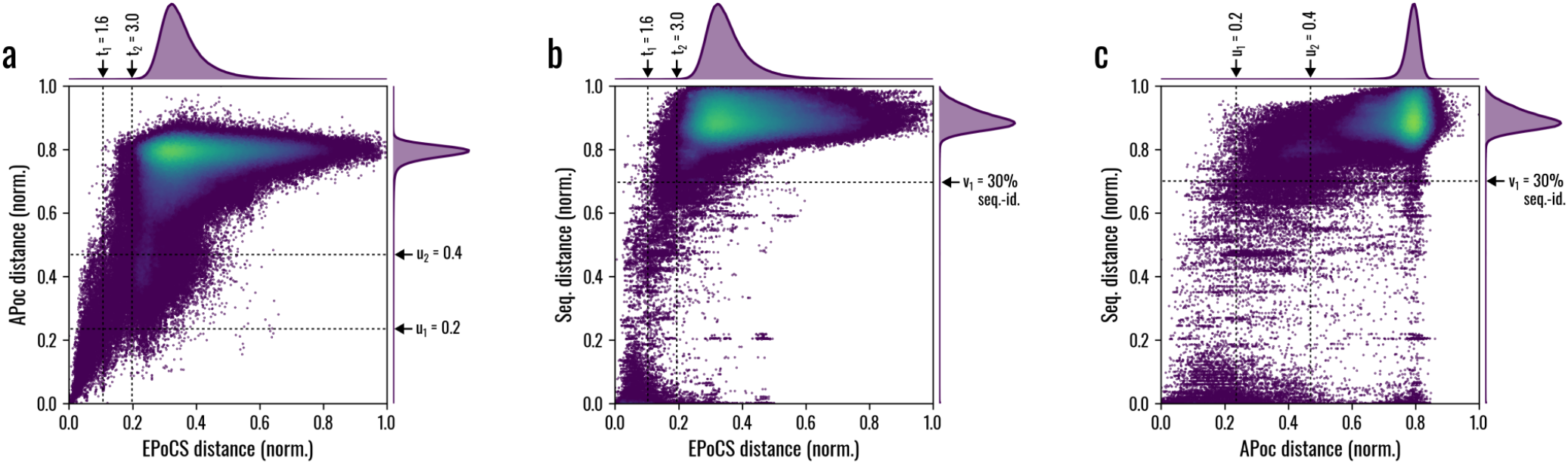
Correlation between EPoCS vs local (APoc) and global (sequence distance) baselines. Pairwise, normalised pocket-pocket distance correlations observed for (a) EPoCS vs APoc, (b) EPoCS vs sequence, (c) APoc vs sequence for a random set of 3500 binding sites from the PDB. EPoCS correlates well with both baseline metrics for pairs that the other measures deem similar. The agreement between the baselines themselves (c), however, proves much noisier – thus pointing to the multiscale character of EPoCS that helps strike a natural balance between local structural, functional and global evolutionary factors. Note that several thresholds *t*_1,2_, *u*_1,2_, *v*_1_ referred to during the analysis (see main text) are labeled with their native (as opposed to min-max normalised) values.

**Table I.** Cluster structure comparison between EPoCS and APoc. Clustering outcomes are compared for two different datasets: (i) a random subset of 3500 pockets from the PDB (PDB-3.5k) and (ii) the intersection of PDB-3.5k with Binding MOAD. Clustering thresholds *t* for EPoCS and APoc are adjusted so as to approximately match the number of induced clusters. The clustering similarity is evaluated using Optimal Transport (OT), and defined as the expectation of the Jaccard index of the best-matching pairs of clusters. The values in brackets are the null similarity index calculated for scrambled clusterings, where cluster sizes were retained, but the pocket labels randomly reshuffled across all clusters. The high agreement between 72% to 86% indicates that the EPoCS metric is locally consistent with alignment-based techniques.

| Subset | EPoCS threshold<br>$t / n\_clusters$ | APoc threshold<br>$t / n\_clusters$ | OT cluster similarity<br>including singletons | OT cluster similarity<br>excluding singletons |
| --- | --- | --- | --- | --- |
| PDB-3.5k | 1.6 / 1917 | 0.2 / 1938 | 0.86 (0.48) | 0.74 (0.29) |
| PDB-3.5k | 2.6 / 1407 | 0.3 / 1435 | 0.80 (0.35) | 0.74 (0.25) |
| PDB-3.5k | 3.0 / 1180 | 0.4 / 1119 | 0.75 (0.27) | 0.75 (0.21) |
| Binding MOAD | 1.6 / 8932 | n/a / 7688 | 0.70 (0.23) | 0.72 (0.18) |

## III. POCKET ATLAS

With EPoCS as similarity kernel, we can use well-established clustering and dimensionality reduction techniques to visualise the metric space and study its behaviour in more detail. The resulting clusterings and visualisations are also of practical interest, for example as a basis for pocket-focused search engines, ligand transfer, selectivity analyses, or (as discussed in section IV) debiasing of train-test splits for validation and benchmarking of ML models.

Here we use distance-based hierarchical clustering (Fig. 1) to group pockets into similarity clusters [33]. Hierarchical clustering is attractive as the agglomeration function allows us to enforce a strict minimum separation between clusters at varying thresholds *t* of the distance measure: i.e., given two distinct clusters *A* and *B*, with pockets *a* ∈ *A* and *b* ∈ *B*, the distance *d*_*ab*_ is at least *t*. Even though the clustering is thus performed on a principled basis, we should keep in mind that the EPoCS metric itself remains a heuristic, and that *t* therefore needs to be selected empirically, with larger values of *t* resulting in fewer but more diverse clusters.

Figure 3 shows the cluster map for a curated set of 96k pockets extracted from 68k PDB structures at an EPoCS threshold of *t* = 1.6 (see *Methods* for details on the dataset curation). The 2D projection was generated using the Mapper algorithm [34], a technique from topological data analysis. Each point on the map corresponds to a cluster of pockets, is scaled according to cluster size, and coloured by functional type (EC class) wherever an unambiguous assignment was possible. The links between clusters indicate the minimum spanning tree as weighted by the minimum distance of approach between the clusters. See the Supplementary Information, Fig. S1, for statistics on the cluster-size distribution and EC-class fragmentation.

**Figure 3.**
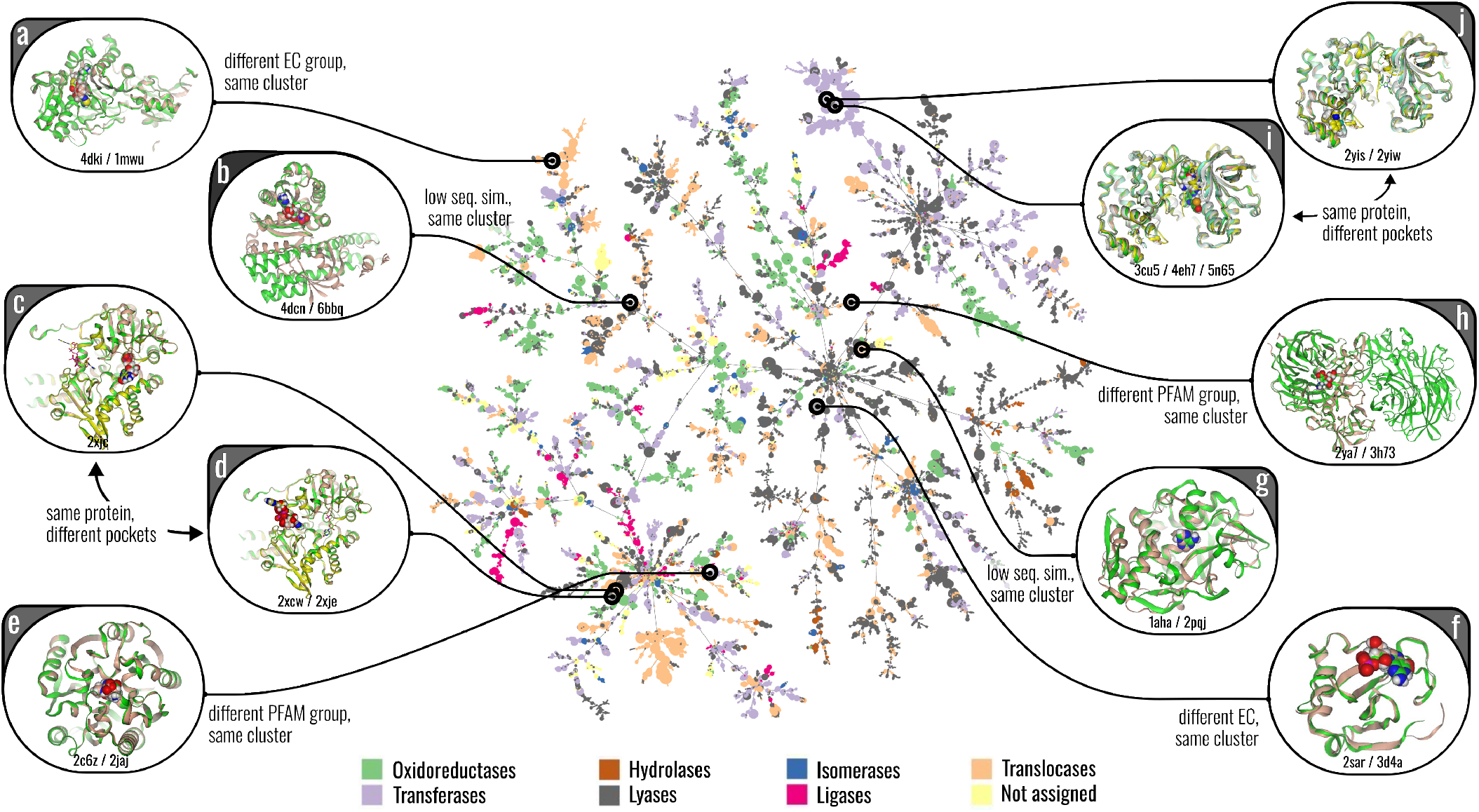
Pocket Atlas. Cluster projection produced by EPoCS for a set of 96k binding sites from the PDB. Each point corresponds to a cluster of pockets, scaled by cluster size. Clusters are coloured according to EC class if all members conform to the same functional label. Detailed inspection reveals that EPoCS reproduces meaningful structural and functional relationships both within and between clusters. Several examples are highlighted to illustrate that: (a, e, f, h) pockets from proteins with different EC or PFAM labels can co-cluster if they are structurally alignable; (c, d, i, j) spatially separated binding sites on the same protein fall into different but nearby clusters; (b, g) sites on proteins with low sequence similarity (< 20%) can cluster together if they are structurally similar. See Table S1 of the Supplementary Information for details on the highlighted complexes.

In the following we provide evidence that this Pocket Atlas organises the pocketome in useful ways: locally, by enforcing structural uniformity within each cluster; regionally, by linking clusters that are biologically or functionally related without an immediate structural homology.

Regarding local structural sensitivity, we have already seen that the EPoCS metric correlates strongly with APoc distances. Applying APoc brute-force to a dataset of this size is, however, not attractive because of the computational cost associated with alignment-based techniques – and furthermore, because of the large number of distant pocket-pocket pairs that fall into a regime where APoc no longer produces any meaningful contrast. We can, however, compare the clustering outcome on a smaller subset of 3500 structures (PDB-3.5k) as introduced in section II, using concepts from Optimal Transport (OT) [35]: i.e., we establish a best-match assignment between the clusters induced by EPoCS to those induced by APoc for clustering thresholds *t* where the total number of clusters is comparable. The expectation of the Jaccard index over the best-matching pairs of clusters is then defined as the OT similarity between the clusterings. The results illustrate (Tab. S1) that EpoCS and APoc have on average a high agreement of 72% to 86% between best-matching cluster pairs, depending on the choice of threshold and exclusion of singleton clusters. This level of agreement indicates that EPoCS locally reproduces alignment-based techniques at a drastically reduced computational cost, while maintaining regional relationships across increasingly structurally decorrelated clusters.

We illustrate this latter point by mapping other chemical and biological information onto the projection. Fig. 4 shows colour maps for: (a) common ligands, (b) drug-likeness (QED) scores and (c) UniProtKB active-site annotations. All three maps display significant local and global structuring, highlighting that EPoCS: (i) aligns well regionally with functional relationships by grouping catalytic sites in consistent ways; (ii) reproduces the well-known fact that similar environments bind similar ligands, but also that some ligands or cofactors (such as ATP) bind to diverse pocket environments – resulting in delocalised hotspots on the map; (iii) captures drug-likeness trends globally, as a likely consequence not just of the different levels of the “true” druggability of the binding-site clusters, but also the intrinsic biases built into the PDB – with some targets and target classes receiving significantly more attention from the drug-discovery community than others.

**Figure 4.**
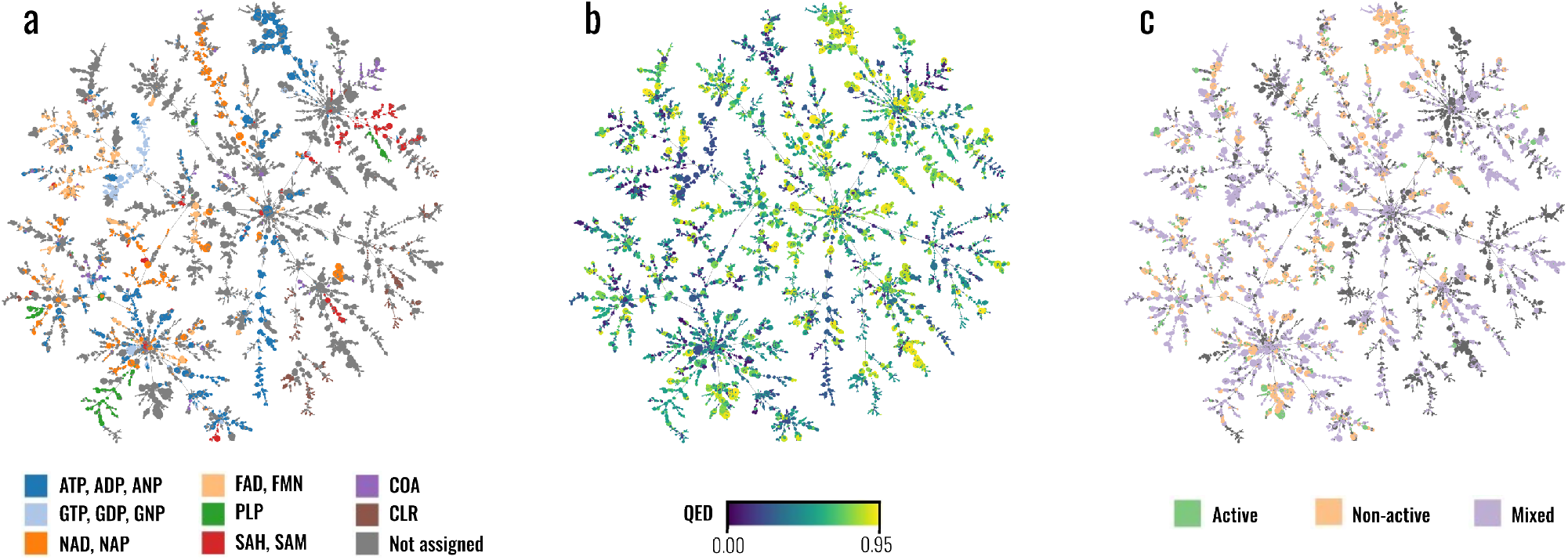
Chemical and functional coherence across EPoCS clusters. EPoCS maintains biochemical relationships locally and regionally. The colour maps indicate: (a) common ligands and cofactors, (b) drug-likeness (QED) scores, (c) active/catalytic site annotations. See *Methods* for details.

Regarding the local cluster structure, one point of concern was that EPoCS could overemphasize the total protein sequence in the local site embeddings, thus potentially clustering together spatially separated sites on the same protein when they are structurally and functionally unrelated. This does not seem to be the case, however, with EPoCS instead allocating these sites to clusters that reside in the same region of the map, but are nevertheless kept well separated by the metric (see examples c, d, i and j in Fig. 3). There are other examples that illustrate that EPoCS seems to conserve many local and regional relationships that are desirable to find in a Pocket Atlas. For example, different EC and PFAM groups can reside in the same cluster if their local structures align well. Even pairs of pockets from proteins with very low sequence identity (< 20%) can fall into the same cluster if they are structurally related. This again highlights that EPoCS maintains high structural sensitivity despite its origin in sequence-based language models.

## IV. POCKET DEBIASING

Estimating the prospective performance of an ML model is hard, and it is not uncommon that models fail dramatically in a real-world setting despite performing well in benchmarks. This disconnect between benchmarking and prospective performance can result not only in a poor model being deployed, but also, potentially, the wrong model: specifically, one that beats the competition on a benchmark and produces good metrics by exploiting accidental bias in the data.

This problem is also pervasive in the development of pocket-centric ML models for protein-ligand docking, affinity or druggability prediction. Here the difficulty in evaluating and comparing diverse methods is partially due to the fact that the PDB (as the data source on which these models are directly or indirectly trained) is a heterogeneous database that was never designed with the training of ML techniques in mind: Its composition is heavily skewed towards certain targets and target classes while containing many ligands/cofactors and/or proteins that are biologically but not therapeutically interesting. Both these biases necessarily persist for random splits of PDB-derived data, as well as the chronological splits that seem to have become an accepted standard.

Sequence-based splits try to improve on this practice, but still fail to prevent accidental data leakage between test and training sets. This is highlighted in Fig. 5, where we used the Pocket Atlas from section III to identify examples in the PoseBusters set (a subset of PDBbind) where chronological and sequence-based splits still resulted in accidental overlap between the PDBbind training and PoseBusters validation set. Such overlap needs to be taken seriously given that ML models are second to none in memorisation, and if there is a simple way to mitigate the issue, then we should make use of it.

**Figure 5.**
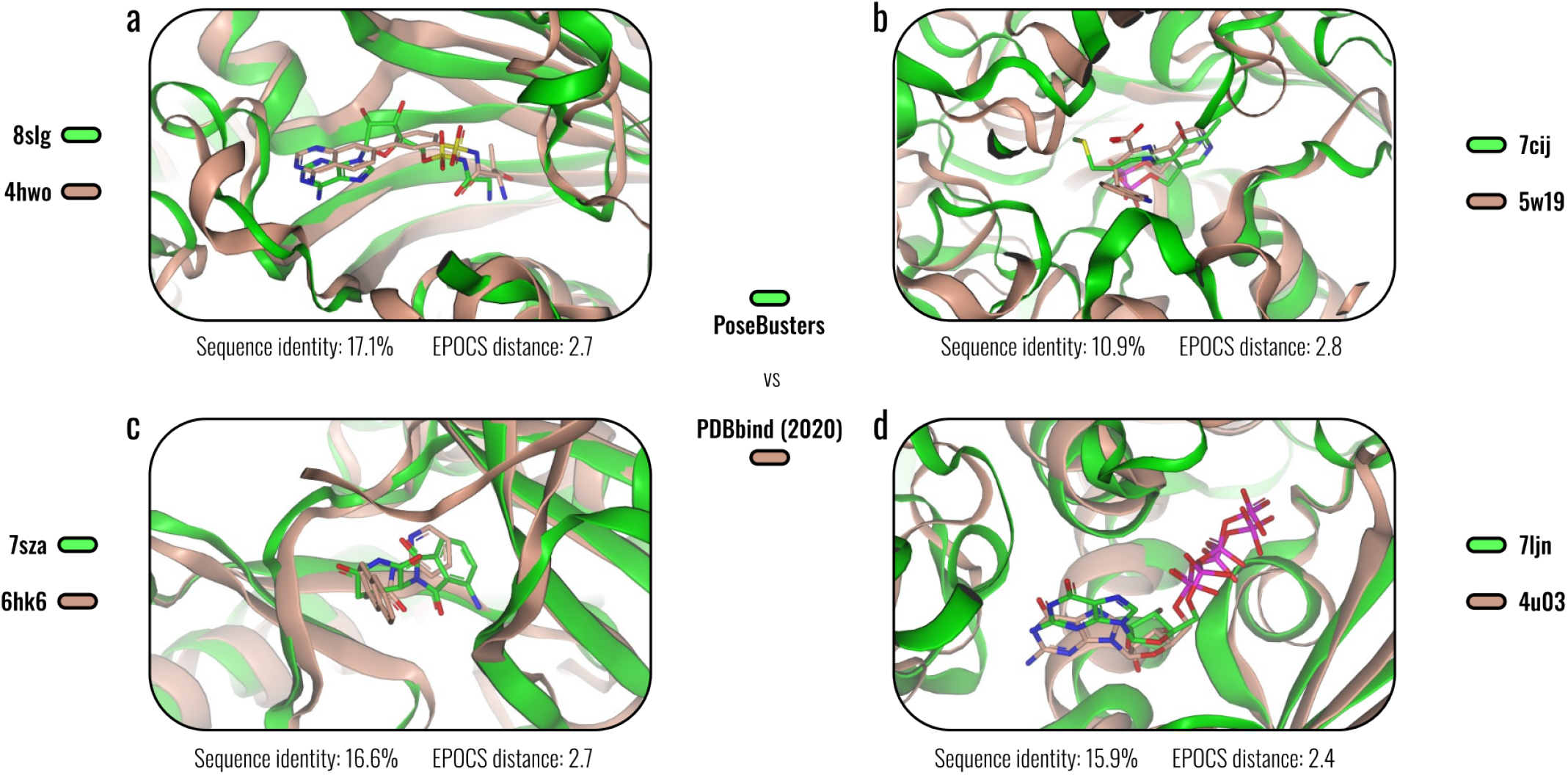
Low-sequence identity, high binding-site similarity. Examples that illustrate accidental overlap between the PoseBusters set and PDBbind 2020 for protein-ligand complex pairs with low sequence (< 20%) but high binding-site similarity. Examples like these can potentially erode benchmarks by rewarding memorisation on seemingly debiased validation sets. We used the EPoCS Pocket Atlas to co-project the low-homology PoseBusters and PDBbind set and thus identify representative examples as shown here. Sequence identities were computed with EMBOSS Needle [36]. See Table S1 of the Supplementary Information for details on the complexes. Note that the complex pairs have an EPoCS distance below *t* = 3.0, which is the largest clustering threshold considered in this work. Appropriate choices for the clustering threshold generally depend on the application, with train-test splits ideally based on larger, and visualisations such as those in Fig. 3 and 4 on smaller thresholds.

Here we show that EPoCS can provide us with one such simple strategy that efficiently decouples train-test splits for pocket-centric ML models. Moreover, we can achieve fine-grained control over the level of debiasing so as to create a sequence of progressively harder train-test splits. Conceptually, there are two different types of splits that can be generated with EPoCS: cluster-based splits, where we adjust the distance threshold *t* and then randomly allocate the resulting clusters to either the train or the test set; and tree-based splits, where we remove entire branches of the minimum spanning tree from Fig. 3 to create a challenging test of a model’s ability to generalise to entirely new pocket and protein contexts.

We illustrate these two debiasing strategies for the training and evaluation of a simple druggability model (referred to as ESM-MLP) that uses ESM-2 embeddings to predict druggability labels on a per-residue basis. Similar to the IF-SitePred model developed by Carbery and co-workers [32], this model is trained on protein-ligand complexes with binary labels *y* ∈ {0, 1} assigned to every residue of every complex depending on whether the residue is contacted by a ligand atom (*y* = 1) or not (*y* = 0) – see *Methods* for details. The cluster- and tree-based splits are constructed such that the training and test sets remain approximately constant in size as the threshold parameter *t* and test branches are varied.

We note that ESM-MLP is performant compared to established druggability baselines including fpocket and PointSite (see *Methods* and the Supplementary Information for details). Here, however, we focus on the properties and behaviour of the splits and split progressions. The areas-under-the-curve (AUCs) for the different training and test sets are summarised in Fig. 6a-c. Note that here we have also added a homology-based split with a 30% sequence identity cutoff for comparison. For the cluster-based splits, we explore three different thresholds *t* = 0, 1.6, 3.0, where 0 corresponds to a fully random split based on singleton clusters. We report test-train averages over three independent instantiations of those splits. For the tree-based splits, we consider three sections *A, B, C* of the pocket tree as test sets, as highlighted in Fig. 6c.

**Figure 6.**
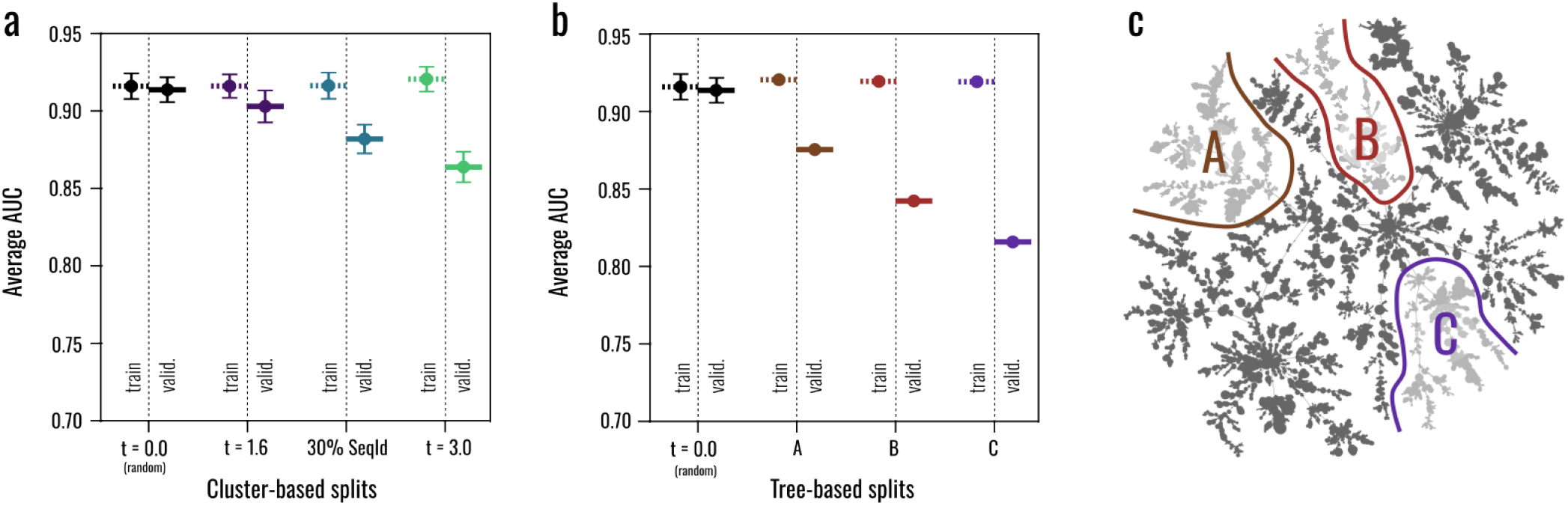
Progression of debiased train-test splits generated using EPoCS. Performance metrics of a pocket-centric ML model (here: for druggability prediction) obtained for different cluster- and tree-based splits generated using the EPoCS metric. Dashed and solid bars indicate training and test set performance, respectively. By varying (a) the threshold parameter *t* for cluster-based splits and (b-c) the test branch for tree-based splits, we obtain a sequence of progressively harder test sets designed to challenge the model, with the classification AUCs for the per-residue druggability scores decaying from > 0.92 for a random split (*t* = 0) to 0.81 for the hardest tree-based split (*C*). A homology-based split with a sequence-identity cutoff of 30% falls in between the cluster-based splits at *t* = 1.6 and 3.0. The error bars indicate the standard error in cases where multiple instantiations of the same split protocol were tested. Note that the absolute sizes of the different train and test sets were kept approximately constant across all splits.

Whereas the AUCs measured for the training sets remain constant at around 0.92, the AUCs obtained for the test sets confirm that we are able to generate successively harder splits, as by design. The easiest tree-based split is about as hard as the hardest cluster-based split. The 30% sequence-identity split, on the other hand, ranks among the easier splits, falling in between two cluster-based splits with thresholds *t* = 1.6 and 3.0 (see our previous discussion around accidental train-test leakage, Fig. 5).

We think that this series of progressively harder train-test splits derived from an expressive similarity metric such as EPoCS are a meaningful but not yet well established practice that can help construct more believable benchmarks. They provide an improved quantitative understanding of how robust and transferable pocket-centric ML models are, whether they are overly reliant on memorisation, or are unstable with respect to training-set selection. It is this type of quantitative insight that helps build trust in the models that are prioritised for further development or deployment.

## V. CONCLUSIONS

Expressive similarity metrics that capture the huge diversity of protein binding sites and their biochemical, functional and evolutionary contexts have many important applications in target and therapeutic discovery. However, striking the right balance between multiple interrelated channels of information at different scales, from local atomic structure to global sequence, is challenging.

Here we have introduced EPoCS, a metric for binding-site similarity based on a state-of-the-art protein language model. The approach shows great promise in capturing relevant, complex patterns at multiple scales, while being computationally efficient. Combined with hierarchical clustering, the metric induces a feature-rich topological projection of the pocketome that we find aligns well with an intuitive understanding of binding-site context from structural, chemogenomic and biochemical perspectives.

Besides enabling interactive queries on pockets of interest for binding-site exploration, EPoCS can serve as a foundation for automatic contextualisation and transfer of ligand information or hit matter across (sub-)pockets and proteins. Building on this work, future research may address to what degree EPoCS-style embeddings can augment dynamical or structural models for druggability and cryptic-site detection.

As deep-learning-based models for pocket-centric tasks such as druggability prediction, co-folding and affinity modelling gain increasing relevance, deep debiasing of train-test splits – as achieved by EPoCS and illustrated for druggability prediction – is urgently needed to construct believable benchmarks that provide fair estimates of how these memorisation-prone methods perform in the real world. These debiased splits not only serve as useful feedback during development of ML models, but help deploy the right model for the right reasons.

## METHODS

### Dataset construction

The cluster maps in Figs. 3 and 4 were derived from a custom subset of the PDB with 96301 structures of liganded binding sites across 68629 PDB entries. The filters that gave rise to this subset are as follows: Ligands classified as additives, stabilizers, saccharides, cofactors, ions or ion clusters were discarded [37], as were PDB structures containing nucleic or non-standard amino acids, and proteins containing chains with more than 1024 residues.

### Site embeddings

Given a protein and a ligand, binding-site residues were determined with Voronoi tessellation, followed by distance-based filtering. While Voronoi tessellation indicates residues with a surface cross-section with the ligand, a radial cutoff of 8 Å between the backbone atoms and any ligand atom is used to remove distant residues.

To obtain PLM residue embeddings, the chain sequences of the PDB structures were extracted and fed as input to a pretrained ESM-2 model (esm2_t36_3B_UR50D) [28]. Given the embeddings for the whole sequence of length *L* (i.e., the *L* × 2560 matrix), a ligand-biased site embedding subsequently used as pocket representation was generated with average pooling over the binding site residues.

### Clustering and visualisation

The distance matrix for hierarchical clustering was generated using the pairwise Euclidean distance between the pocket representations. The linkage policy for the clustering was based on the minimum distance of approach between any of the members of a pair of clusters. To avoid redundancy and reduce the memory footprint, pre-clustering was applied with a small distance cutoff (*t* = 1.0) and one representative was selected from each cluster as a clustering landmark.

For the pocket maps in Figs. 3 and 4, clusters were merged up to a distance threshold of *t* = 1.6. We note that the pre-clustering step can result in violations of the distance guarantee between clusters, which is why the final distance between nearby pairs of clusters was re-evaluated to construct the effective cluster-cluster distance matrix. Edges of the Pocket Atlas were calculated from the cluster-distance matrix using a minimum spanning tree algorithm as implemented in the tmap library [34]. The interactive version of the map was constructed using the Faerun package [38].

### Comparison to APoc and SeqId

We evaluated the correlation of EPoC distances with APoc [17, 39] and a simple sequence distance measure over a random subset of ∼ 3.5k pockets. We used the same binding-site definition for APoc and EPoCS as defined above. For each pocket pair (*a, b*), Apoc similarity scores were calculated in both directions, *a* vs *b* and *b* vs *a*, to account for differences in normalisation, with the average of the two scores used as the final similarity *s*. We used *d* = 1 − *s* as the associated distance value for both APoc and sequence similarity.

### Clustering distance

We used optimal transport over the cluster-cluster Jaccard index to measure the similarity between the clusterings induced by EPoCS vs APoc (Table S1). We used identical clustering procedures for both the EPoCS and APoc metrics (see clustering details above). The distance thresholds for both techniques were chosen so as to approximately match the resulting number of clusters (for example, we match the APoc threshold *u* = 0.2 to an EPoCS threshold of *t* = 1.6, resulting in, respectively, 1938 vs 1916 clusters). To additionally compare to the binding-site families defined by Binding MOAD [40], which are ultimately derived from the APoc metric, we filtered the set of PDB IDs down to those present in both sets. This latter comparison was based on PDB IDs (not pockets) in line with Binding MOADs description of site families.

We calculated the Jaccard index for all pairs of clusters induced by the different metrics (i.e., the number of pockets shared by the cluster pair over the total number of pockets associated with that pair). Optimal transport [35] was applied to the cluster-cluster similarity matrix to find the best-matching cluster assignment. We use the average of the Jaccard index for the assigned clusters as similarity for the two clusterings. As a null baseline, we also report the final similarity obtained for scrambled clusterings, where the number and size distribution of the clusters is retained, but cluster membership is shuffled.

### Map annotation

We used the SIFTS database [41, 42] to annotate the PDB structures with their Enzyme Commission (EC) numbers. A cluster of binding sites was labeled as belonging to a certain EC class if: at least one member of the cluster belongs to that EC class; and there is no pocket in that cluster belonging to any other class.

Active-site information was retrieved using EBI’s API. Residue mappings from UniProt to the PDB were extracted from SIFTS. For a given PDB ID and EPoCS binding site, the pocket was labeled as ‘active’ if the mapped residue ID was among the binding-site residues.

For the annotation of the Pocket Atlas, a cluster was labeled as ‘active’ if all pockets included in that cluster were ‘active’. For some structures, active site information is missing or not available. Clusters of pockets that are associated with activity labels but that do not incorporate the ‘active’ residues themselves, are labeled as ‘non-active’. Other clusters are grouped under the label ‘mixed’.

To identify clusters with diverse sequences, pre-generated sequence-based clusters with 30% identity cutoff based on MMseqs2 [43]) were adopted from the PDB. For EPoCS clusters split across several sequence clusters, pairwise sequence alignments were generated and their pairwise sequence identities calculated. Pocket clusters associated with at least one pair of proteins with sequence identity less than 30% were labeled as sequence ‘diverse’.

QED scores were calculated with RDKit [44], with the maximum QED score used to represent the cluster.

### Druggability modelling

The ESM-based druggability model MLP, consists of a multilayer perceptron (MLP) with residue ESM-embeddings as inputs and a single log-probability channel as output. The MLP includes a single hidden layer with 12 nodes. Binary training labels for residues were assigned based on the same Voronoi tesselation procedure as used by EPoCS. We used an Adam optimizer with a constant learning rate of 0.001 and weight decay of 0.001. The MLP’s dropout rate was set to 0.3. The models were trained with a log-likelihood loss function on the ‘general’ set of PDBbind 2020.

### Comparison to fpocket and PointSite

Despite its simplicity, ESM-MLP is found to be performant – see Figs. S2 and S3 for a comparison to fpocket and PointSite. We note, however, that training set consistency is not maintained across the three models. Hence, the structures which are present in both HOLO4K (one of the test sets of PointSite) and the ‘general’ PDBBind set were selected as an additional test set that allows for a direct comparison between PointSite and ESM-MLP.

Unlike the ESM-based model, fpocket does not assign continuous druggability scores to individual residues. We therefore used membership labels of the residues belonging to a predicted binding site as a binary proxy for an fpocket druggability score. PointSite, on the other hand, produces atomic druggability scores, which we translated into residue-level scores using max pooling. For PointSite, fpocket, and ESM-MLP, cluster AUCs were evaluated by concatenating the modelled residue scores and calculating the true-positive and false-positive rates over the ligand-based ground-truth contact labels as a function of score threshold.

### Software

We used pdb-tools [45], pymol [46] and biopandas [47] to process PDB structures, and RDKit for molecular processing. Hierarchical clustering was performed using SciPy [48]. Optimal transport metrics were evaluated using the POT package [35].

## Data availability

The PDB subsets and clusterings are available publicy at https://github.com/tugceoruc/epocs/tree/pre-release/data.

## Code availability

Code for the generation of site embeddings, site projection and visualisation is available at https://github.com/tugceoruc/epocs.

## Acknowledgements

We are grateful to Chris Murray and Lisa Ronan for thoughtful comments on the manuscript. We thank Lucian Chan and Fred Ludlow for fruitful discussions.

## Author contributions

TO: Design, implementation, execution, analysis. MK: Code, discussions. TD and MV: Discussions, analysis. CP: Design, analysis, supervision.

## SUPPLEMENTARY INFORMATION

**Figure S1.**
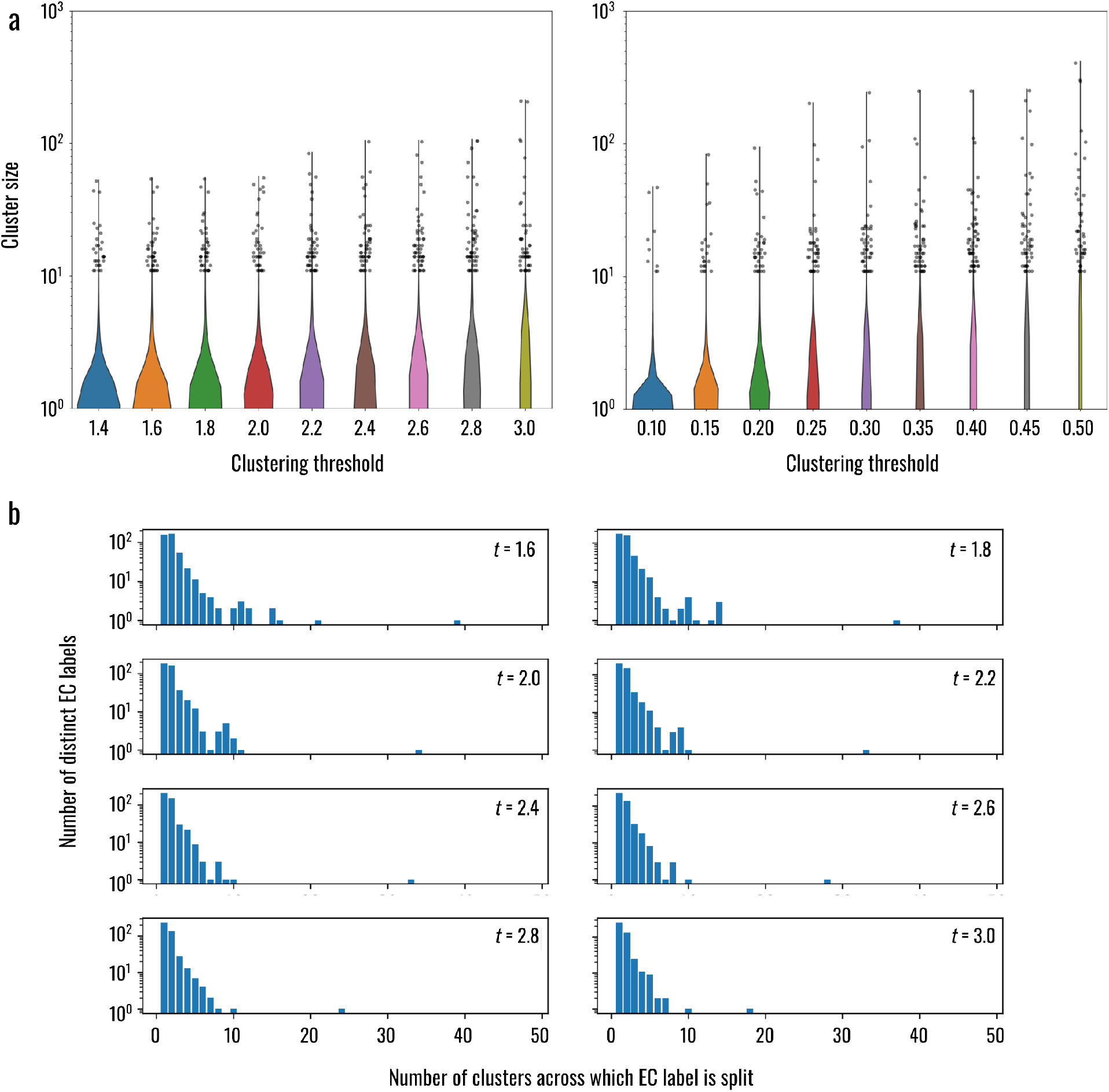
Clustering statistics at different separation thresholds. (a) Violin plots of the cluster size distribution induced by (left) EPoCS and (right) APoc for several clustering thresholds. (b) Degree of fragmentation of EC groups across EPoCS clusters at different clustering thresholds. The vast majority of EC groups are contained within fewer than 10 clusters, indicating that the clustering outcome is well aligned with functional annotations over the range of thresholds explored here.

**Figure S2.**
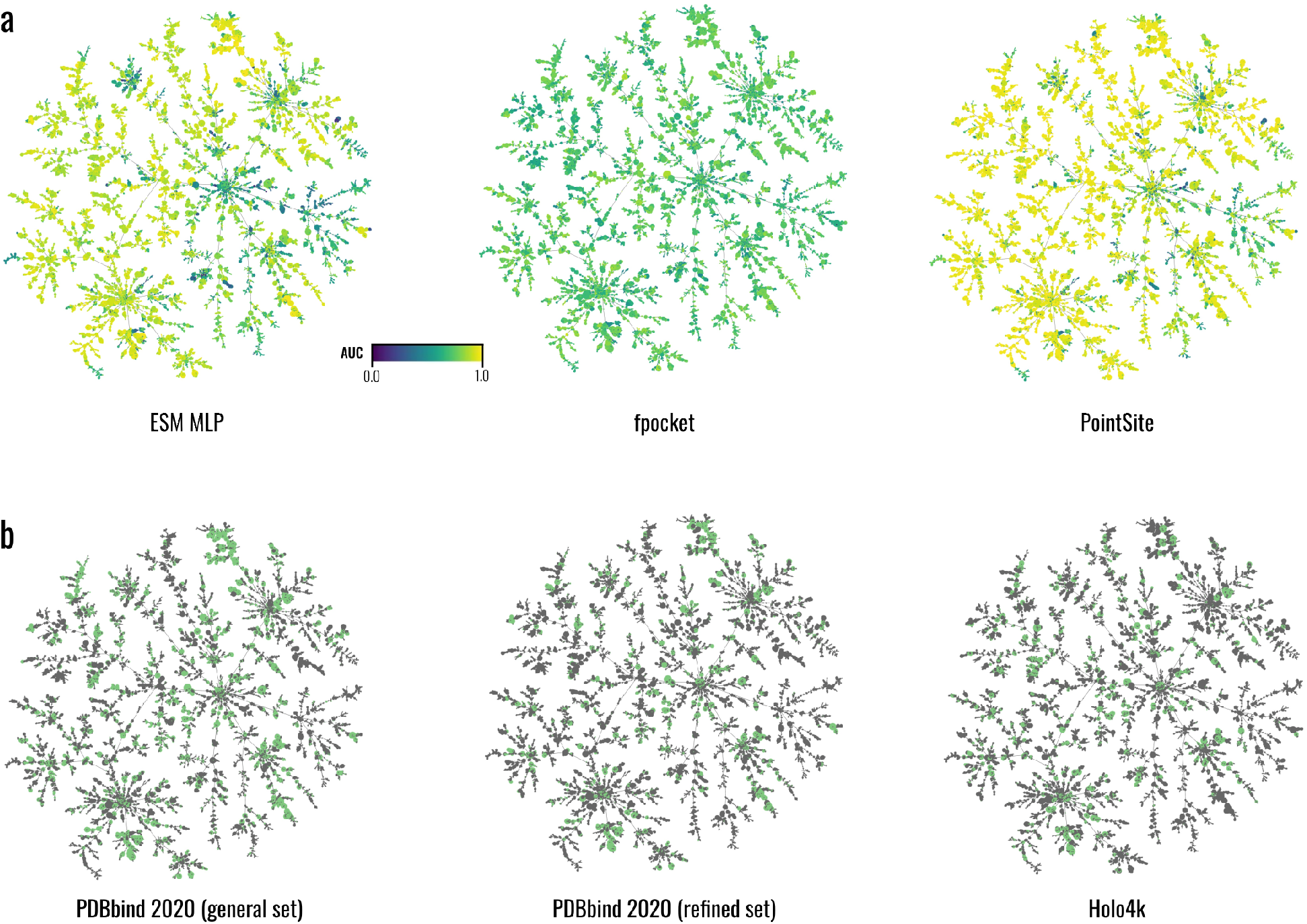
Comparison of druggability prediction performance across the EPoCS atlas. (a) Per-cluster AUCs assessing the predicted per-residue druggability scores generated by the ESM MLP, fpocket and PointSite vs the ground-truth contact labels. Despite its simplicity, the ESM MLP rivals PointSite in the quality of its predictions, with fpocket-based proxy scores falling behind the other two methods. Interestingly, the ESM MLP and PointSite are broadly aligned in how their AUCs vary across the pocket atlas, indicating a similar concept of druggability learnt by these two models. These global trends are less perceptible in the case of fpocket, which is a volume-based rather than contact-based approach. (b) Distribution of PDBbind 2020 subsets used in the training and evaluation of the ESM MLP across the pocket atlas.

**Figure S3.**
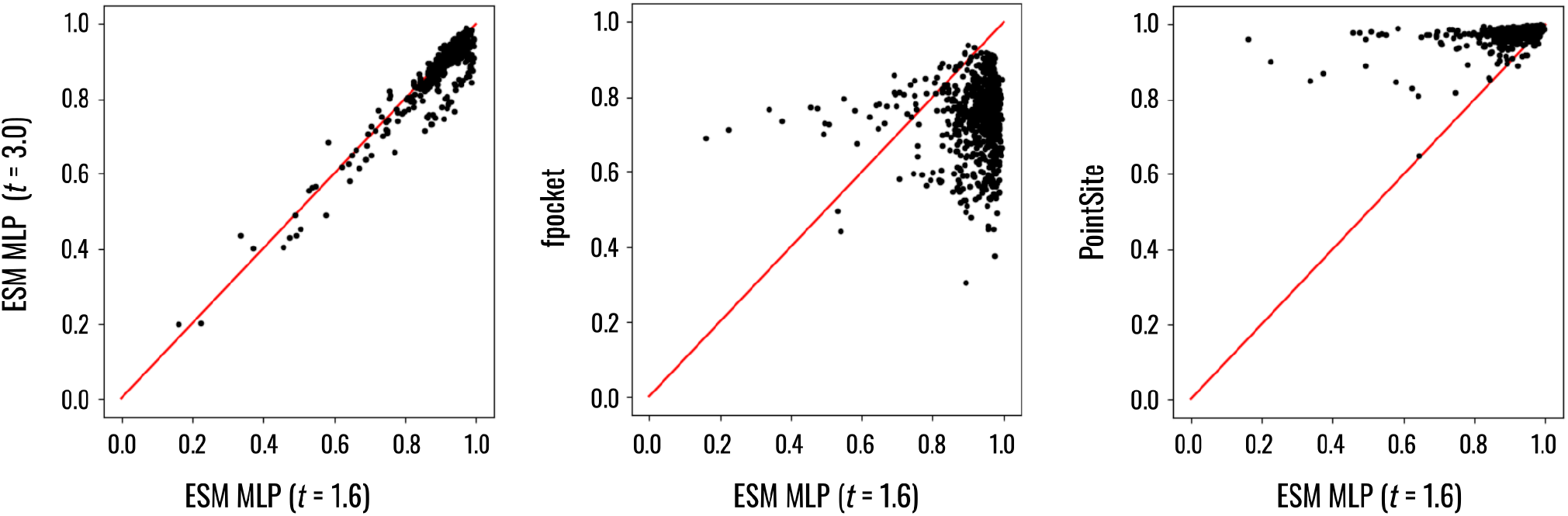
Score correlations on the intersection of HOLO4K and PDBbind. Score correlations among four druggability models (ESM MLP trained on sets derived from two different EPoCS thresholds *t* = 1.6, 3.0, fpocket and PointSite. HOLO4K is part of PointSite’s test set, making this a fair comparison between the ESM MLP models and PointSite.

**Table S1.**
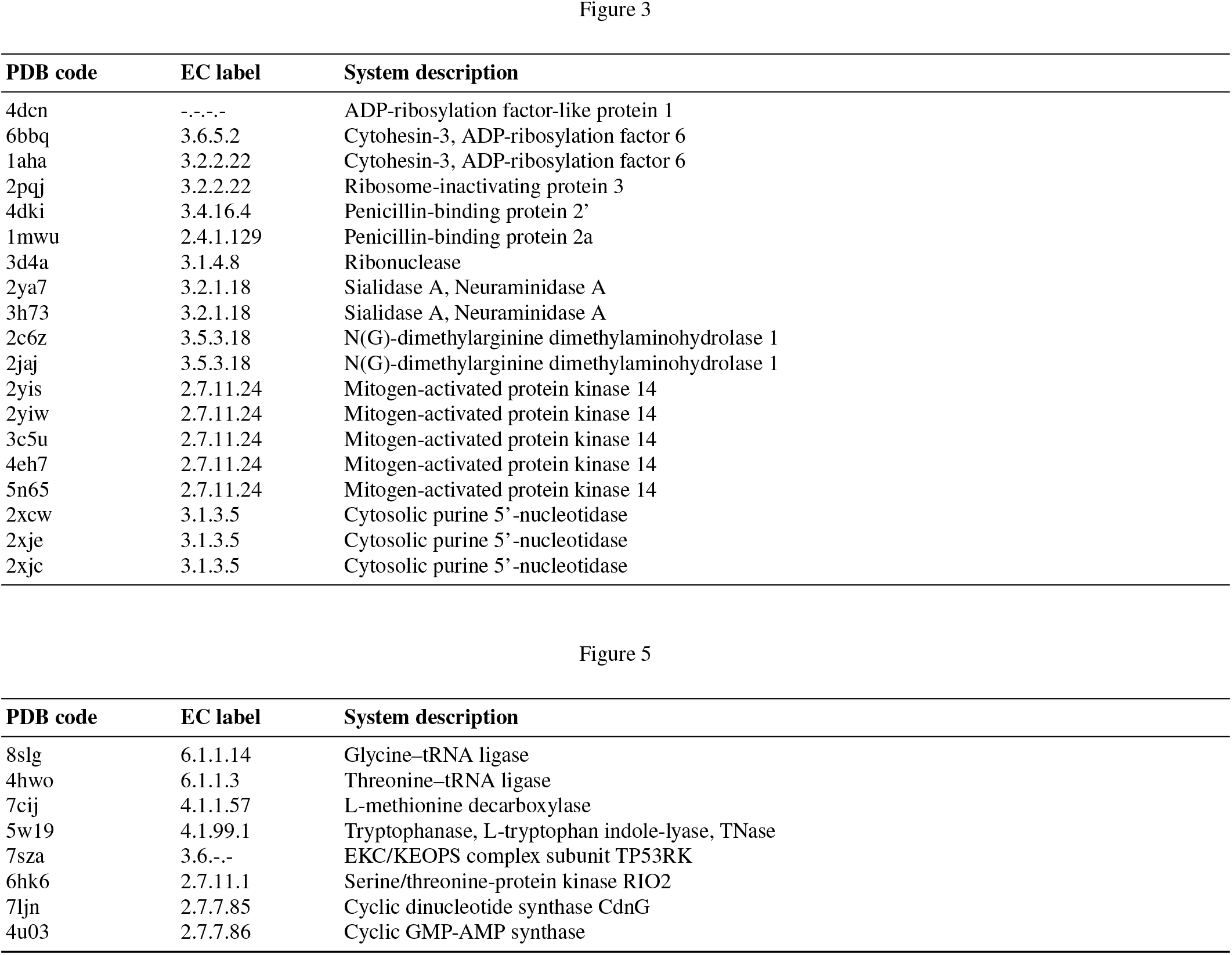
PDB codes referenced in Figs. 3 and 5 of the main text, with EC label and system description.

